# Jupytope: Computational extraction of structural properties of viral epitopes

**DOI:** 10.1101/2022.03.22.484725

**Authors:** Shamima Rashid, Ng Teng Ann, Kwoh Chee Keong

## Abstract

Epitope residues located on viral surface proteins are of immense interest in immunology and related applications such as vaccine development, disease diagnosis and drug design. Most tools rely on sequence based statistical comparisons, such as information entropy of residue positions in aligned columns to infer location and properties of epitope sites. To facilitate cross-structural comparisons of epitopes on viral surface proteins, a python-based extraction tool implemented with Jupyter notebook is presented (Jupytope). Given a viral antigen structure of interest, a list of known epitope sites and a reference structure, the corresponding epitope structural properties can quickly be obtained. The tool integrates biopython modules for commonly used software such as NACCESS, DSSP as well as residue depth and outputs a list of structure derived properties such as dihedral angles, solvent accessibility, residue depth and secondary structure that can be saved in several convenient data formats. To ensure correct spatial alignment, Jupytope takes a list of given epitope sites and their corresponding reference structure and aligns them before extracting the desired properties. Examples are demonstrated for epitopes of Influenza and SARS-CoV2 viral strains. The extracted properties assist detection of two Influenza subtypes and show potential in distinguishing between four major clades of SARS-CoV2, as compared with randomized labels. The tool will facilitate analytical and predictive works on viral epitopes through the extracted structural information.

**Key Messages:**

- Jupytope combines existing 3D-structural software to extract the properties of viral epitopes into a convenient text or csv file format
- The structural properties serve as parameters or features that quantitatively capture viral epitopes
- Association of structural properties to viral subtypes (for Influenza) or clades (SARS-CoV2) is demonstrated with a simple XGBoost model
- Structure datasets mapped to SARS-CoV2 WHO clades and Pango lineages, as well as chain annotations are available for download

## Introduction

Viral membrane proteins, such as SARS-CoV2 Spike and Haemgglutinin (HA) of Influenza play a key role in host attachment, infection and immune response.

### SARS-CoV2, Influenza Pandemics and the Roles of Spike and Haemagglutinin in viral mechanisms

Severe Acute Respiratory Syndrome Coronavirus 2 (SARS-CoV2) is a virus that triggered the novel Coronavirus (COVID-19) disease. It was first identified in November 2019, in Wuhan, the capital of China’s Hubei province (17). Efforts to contain it there were unsuccessful, and since then, it has spread globally, resulting in World Health Organization (WHO) declaring the large-scale outbreak a pandemic on March 11, 2020 (1). As of Feb 25, 2022, WHO reported 430,257,564 confirmed cases, including 5,922,047 deaths globally (29).

Influenza A virus (IAV) is highly contagious and has continually posed significant danger to global human health and life (33). Emergence of recombinant pathogenic Influenza A virus (IAV) strains are unexpected events with rapid spread leading to occurrence of pandemics (26), such as the 1918 Spanish flu (H1N1), 1968 Hong Kong flu (H3N2) and 2009 Swine-Origin flue (H1N1). These pandemics respectively resulted in 20 to 50 million (14), one to four million (25) and 151, 700 to 575,400 deaths (32) in the human population worldwide. Pandemics often cause increased morbidity with vast clinical and economic impact (26).

SARS-CoV2 and IAV primarily transmit between humans via aerosolized respiratory droplets and airborne particles containing the virus (17; 3). It predominantly affects the respiratory system (23), with a broad spectrum of severity ranging from mild common symptoms including fever, cough, shortness of breath and fatigue (1; 40) to severe acute respiratory illnesses that may be life-threatening (40; 10). Furthermore, a SARS-CoV2 carrier can be asymptomatic and spread the disease unknowingly (17). SARS-CoV2 is a positive-sense single stranded Ribonucleic Acid (RNA) genome (2), belonging to the Betacoronavirus family (11). Genome sequence analysis revealed resemblance of the virus to SARS-CoV1 (approximately 79%) and Middle East respiratory syndrome coronavirus (MERS-CoV) (approximately 50%) (23; 22).

Spike protein on the surface of the virus is a crucial factor mediating viral entry and infection by receptor recognition, cell attachment, and fusion. It is a trimeric class I TM glycoprotein, present in all kinds of Human coronaviruses (HCoVs), and Hemagglutinin (HA) of Influenza belongs to the same class (11). The major role of spike protein in viral infection suggests that it is a potential primary focus target for most vaccine design and development efforts (15). SARS-CoV2 spike protein consists a total of 1273 amino acid residues. The S1 subunit (residues 14 to 685), and the S2 subunit (residues 686 to 1273) are responsible for receptor binding and membrane fusion, respectively (11). Entry of virus into the body is enabled through receptor binding of the viral spike protein by recognizing the Angiotensin-converting enzyme 2 (ACE-2) receptors on the surface of the human alveolar epithelial type II cells (11). Upon entry, a single positive strand of RNA is released into the cytoplasm of the host cell (2). Viral fusion activity is activated under low pH, due to the formation of endosomes, promoted as a result of the binding of S1 subunit with ACE-2 (11).

IAV is a characterized by an enveloped negative-strand RNA virus, belonging to the Orthomyxovirida family. Its genome comprises of eight RNA segments, with segments four and six encoding for key surface glycoproteins, HA and Neuraminidase (NA) respectively. In accordance with its antigenicity, IAV can be divided into 18 HA (H1-H18) and 11 NA (N1-N11) subtypes. The strain was determined by the combination between different HA and NA subtypes (40). The HA of influenza virus plays a vital role in the early stage of virus infection as it is responsible for binding of the virus to cell surface receptors, and subsequently, mediating the release of the viral genome into the cytoplasm of the host cell through membrane fusion. The necessary component of the receptor for IAV was believed to be the sialic acid. IAV recognizes N-acetylneuraminic acid as the receptor. A number of amino acid residues constituting functional domains, namely, receptor binding site and fusion peptide are revealed to be conserved among the HAs across IAV subtypes, located on the HA1 and HA2 subunits respectively (24). HA1 contains the receptor binding domain, with 190-helix and 220-loop. HA1 harbours many epitope residues, while the HA2 subunit mediates membrane fusion and viral entry (4).

### Viral Epitopes Characterization and Relation to Clade or Subtype

Current clade, lineage or subtype nomenclature describes genetic differences between virus strains and is based on phylogenetic relationships between ancestral and descendant sequences. The major known clade groups of SARS-CoV2 correspond to G, L, S and V, with all of the variants of concern (VOC) belonging to clade G, while Clade L was shown to be characteristic of the early strain in Wuhan (39). Additionally, Group O comprises lineages that are ‘others’ which cannot be clearly linked to the 4 major clades.

However, the association of clades with antigenic properties or phenotypic outcomes is unclear. Recently, a study linking SARS-CoV2 clades with patient disease outcomes was published (39). They concluded that L and V subtypes were more strongly associated with severe forms of the disease such as hypoxia, requiring oxygen supplementation. Epitopes of Spike and HA, play a significant role in disease outcomes (e,g, in vaccine selection or natural antibody response), but their connection to SARS-CoV2 clades or Influenza subtypes, is less clear.

Epitopes of viral antigens may be classified as continuous (linear in sequence) or discontinuous (non-contiguous in sequence) besides others such as neotopes or mimotopes. Discontinuous epitopes, are defined by the spatial orientation of particular residues brought into proximity by folding of the viral polypeptide chain. It is generally difficult predict epitopes, precisely because they are defined in response to antibodies and are not intrinsic features of the antigen (38). Structure based methods for epitope prediction, use physico-chemical descriptors as features, such as the Half Sphere Solvent Exposure (HSE) (9) used by DiscoTope-2.0 (19) and BEPro (36). Other descriptors are the residue depth provided by MSMS (31), solvent accessibility scores of NACCESS (12), and secondary structures defined according to DSSP (16). Rubinstein et.al., included up to 45 structure derived properties for epitopes prediction (30). However, most prediction methods are “underused and characterized by a high rate of false positives”, according to Kozlova, et.al., (18). A good review of epitope prediction available in (19).

The use of structure-derived features for epitope prediction has proven significant over the use of sequence-profile based methods for epitope prediction, (19). It is motivating to present a work that enables the easy extraction of structure based features, which would drive the generation of structure-derived epitope datasets. We hypothesized that the epitope’s structure data could be associated with their viral clade or subtype. Such an association indicates that the antigenic profile has been sufficiently captured, and it would be possible to infer viral group associations from the epitopes.

This work presents a python based tool to automatically extract structure-derived properties of viral epitopes of SARS-CoV2 and Influenza A. The tool combines established properties such as solvent accessibility and HSE into a single convenient output, from the structure of a given viral antigen and a reference. Dependent software or python libraries need only be installed once and an initial download of structure files is required..The program will extract the associated data needed for a given list of PDB files. Further, Jupytope’s output is verified by checking if virus clade or subtype specific associations can be recovered from the extracted data, using a simple XGBoost model compared to a random baseline. In the rest of the manuscript, the phrase ‘virus group’ is collectively used to refer to SARS-CoV2 clades and Influenza subtypes.

It is proposed that the tool facilitates the comparison of epitope characteristics across different structures and will aide structural bioinformatics workflows by preventing messy “cut and paste” from individual programs. Further, the extracted structure-derived epitope features can be a good starting point for machine learning algorithms. For instance, it is suited for deep learning of data representations from the features. One application is the prediction of epitope vs non-epitope residues, where the extracted data serve as a large pool of positive labelled epitope residues that are (i) accurately aligned with respect to their reference structures and (ii) labelled with their respective virus groups. This work also presents SARS-CoV2 PDB structures that have been annotated with the respective Spike chain monomers, their GenBank Identifiers (IDs) and corresponding Clade and Pango lineages (27), to facilitate further analysis and predictive works.

The manuscript is organized as follows. Methods describes the development of the SARS-CoV2 and Influenza structure datasets, the program overview, and the epitopes used in its construction. Results and discussion describe the structure-derived epitope features, and their performance with a simple XGBoost classifier in recovering virus groups. The conclusion summarizes the findings and suggests the applications of the features. The Appendix ‘List of Extracted Structural Properties’ lists the 25 features and the source programs used to generate them.

## Methods

Data pre-processing and labelling was carried out before extracting the structural information using Jupytope.

### Datasets

#### Influenza A

Influenza A records for H1N1 and H3N2 were downloaded in July 2018 from the Influenza Research Database (IRD) using their 3D-protein structure query (41). 58 H1N1 and 72 H3N2 records containing structures and their respective strains were retrieved. The structure files were downloaded from the Research Collaboratory for Structural Bioinformatics (RCSB) Protein DataBank (PDB)(5) and manually annotated as follows. Structures containing incomplete or missing HA1 segments (e.g. fragments less than 30 residues long) were removed. The HA chain in each structure file was identified and the oligomer unit (trimer, monomer, etc) was recorded. Antibody chains (if present) and bound receptors were also recorded. Complexes containing sialic acids or N-acetylglucosamine (NAG) were collectively included as having receptors. All of the structures were determined by X-Ray crystallography. The final set contains 97 (43 for H1N1 and 54 belonging to H3N2, respectively) annotated Influenza structures, given in the Supplement.

#### SARS-CoV2

The reference Spike protein sequence was first downloaded from GISAID (28). hCoV-19/Wuhan/WIV04/2019 (WIV04) with accession: EPI_ISL_402124, was responsible for the first known outbreak in Wuhan. The RCSB PDB was queried in January 2022 using this reference sequence with an E-value of 0.1 and an identity cut-off of 70%, resulting in 686 polymer entities. After removing structures shorter than 900 residues or extremely large files, a total of 398 structure files were downloaded.

However, not all the structures could be utilized. Owing to the difficulty of obtaining complete crystals for transmembrane proteins, there is often a discrepancy between the number of residues stated in the PDB sequence records (referred here as SEQRES) and the number available in atomic coordinates (referred here as SEQATOM). After further checks and selecting for structures having SEQATOM length of at least 700, the final set contained 353 PDB structures, all of which were derived from electron microscopy.

Figure 2 shows the resolution of X-ray diffraction (Influenza) and cryo-EM (SARS-CoV2) crystals in this work. Typically, a resolution of ≤ 2.5*Å* is considered acceptable quality in structure datasets (13; 34) and ≤ 1.5*Å* is considered sufficient atomic detail for drug or ligand placement studies. Here, owing to the difficulty of obtaining sufficient numbers crystal structures with complete domains of HA and Spike proteins, we did not disregard structures based on their resolution.

**Fig. 1.**
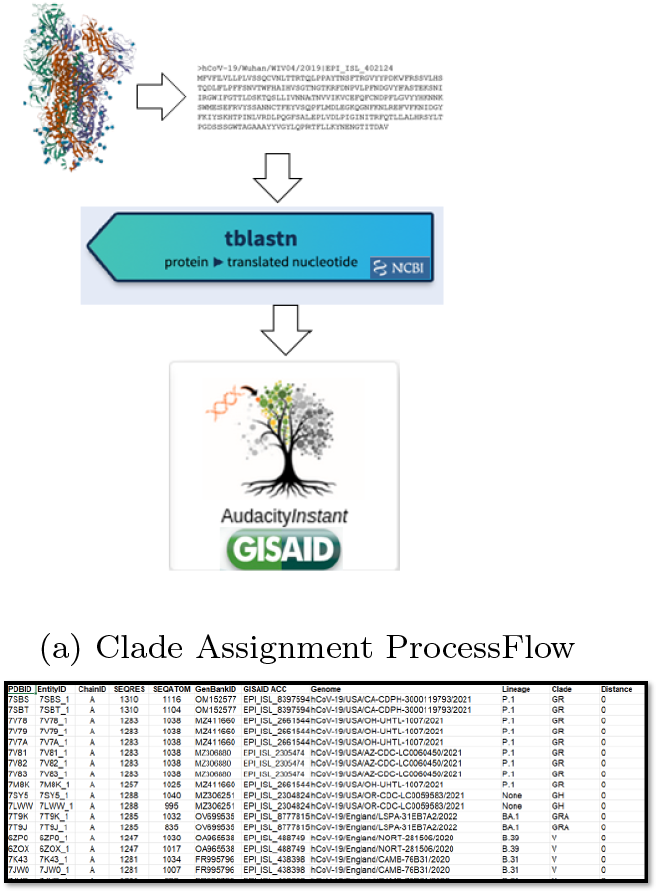
SARS-CoV2 Clade Assignment

**Fig. 2.**
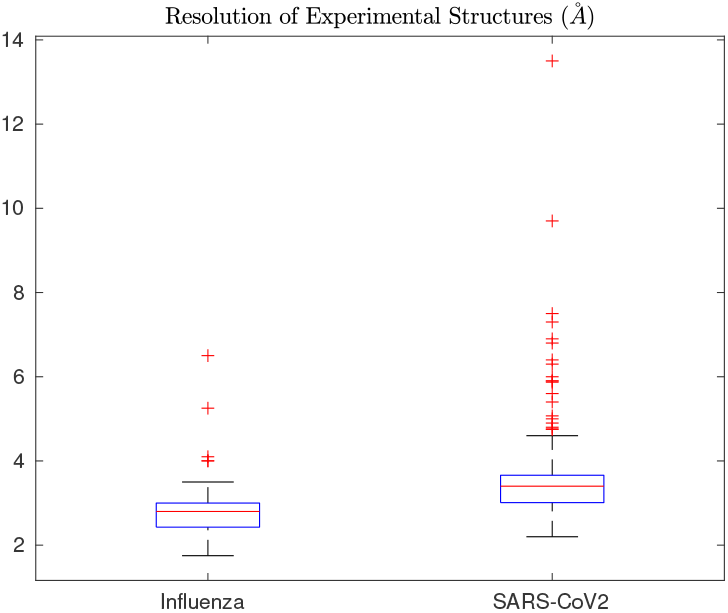
Resolution of Structures. Boxplots shown are for Influenza (X-ray) and SARS-CoV2 (cryo-EM) crystals that were used to extract properties of epitopes. A total of 97 and 353 structures were used for Influenza and SARS-CoV2, respectively.

The PDB structures were not labeled directly with clade or lineage information and existing 3D SARS-CoV2 databases have either too few records, or lack the clade or lineages (unlike for Influenza). Hence, to address this gap and to investigate the antigenic associations between viral epitopes and viral groups, the downloaded structures were assigned clades. The workflow represented in Figure 1(a) shows our clade assignment scheme. SEQRES of PDB structures were queried using the tblastn software of NCBI (6) to retrieve the complete genome nucleotide record (Genbank ID (7)) of the closest matching strain (almost 30,000 bp), against the Betacoronavirus database. The closest hits yielded E-values of 0. The genomic nucleotide sequences were individually queried against GISAID’s Audacity web application and the top matching clade, lineage and corresponding distances were recorded. Figure 1(b) shows a sample of the annotated Genbank IDs and GISAID clades, assigned to the 353 SARS-CoV2 structure files. The Pango lineages were also recorded. The full file, is available in the Supplement and can be applied as a dataset for inferring or characterizing SARS-CoV2 clades or lineages from structure properties.

### Epitope Numbering Scheme

A list of epitope sites located on HA and Spike structure surfaces were collected from the literature, and aligned with reference structures to preserve numbering. The numbering scheme and reference structure used for each type of virus are provided in Table 1.

**Table 1.** Source of Epitope Sites

| Virus | Epitope Sites | Total Residues | Ref. Struct |
| --- | --- | --- | --- |
| Influenza (H1N1) | A-E (8) | 161 | 1RU7 |
| Influenza (H3N2) | A-E (21) | 241 | 1HGD |
| SARS-CoV2 | E1-E9 (35) | 228 | 7DZW |

Owing to the small number of experimental structures available especially in comparison to the large amount of genomic sequence data, only single chain structures were considered here, which allowed the use of monomers in the analysis. Figure 3 shows the epitopes mapped onto reference PDB structure for H1N1, recreated from (8).

**Fig. 3.**
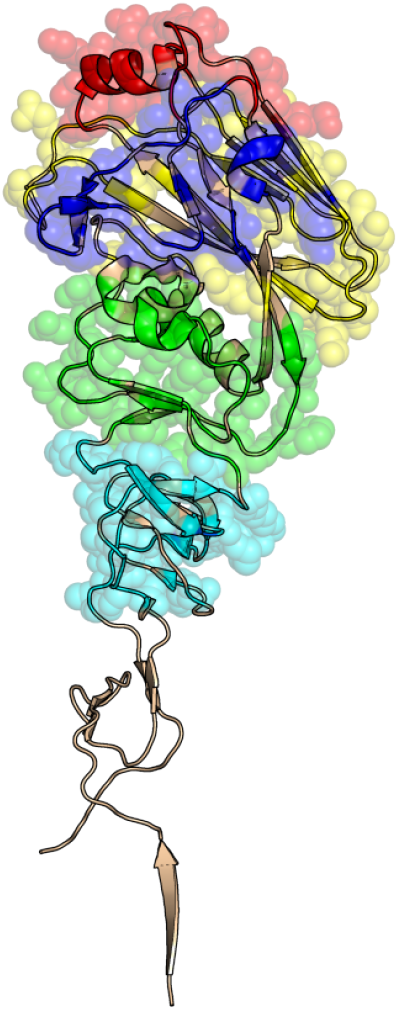
Visualization of Epitope Sites A-E for H1N1, using reference structure with PDB ID 1RU7, Chain A. Figure recreated from (8). Site A is colored in blue, Site B in red, Site C in cyan, Site D in yellow and Site E in green.

Single HA chains were extracted from the collected datasets using ChainSplitter code written by David Cain in 2012. The structural properties of their epitope residues were extracted according to the reference structures and numbering scheme as in Table 1.

### Program

Jupytope is available for download from the github repository https://github.com/shamimarashid/Jupytope. The code was developed in Jupyter Lab Notebook version 2.2.8, using a python 3.6.11 interpreter. The program relies on the Biopython packages (Bio.PDB, Bio.AlignIO) and other structure processing and parser libraries for individual programs listed as follows: NACCESS, DSSP, ResidueDepth (MSMS) and Half Sphere Solvent Exposure (HSExposure). The Kyte and Doolittle hydrophobicity scale is used to assign values to the epitope residues (20). Further, the normalized B-factor (B Norm) was calculated according to the protocol in (37), defined in Jupytope as function. To compute B Norm, the average B-value for all atoms for a single residue is normalized using the average of all residues within the HA chain. B-Norm values are not used in the features analysis for Spike, since they are not meaningful for cryo-EM crystals. A full list of dependencies are available within the notebook or README file at the above github repository.

Figure 4 shows the schematic of information flow in Juyptope. The inputs to the program are a (i) file containing a list of PDB ID and chains, (ii) a directory containing the PDB structures, (iii) a reference structure and (iv) a numbered list of epitope residues of interest (*S*). The output of the program is an extracted list of structural properties of epitope residues (*F*), corresponding to the residue properties calculated by individual programs. These are consolidated and saved as a text or csv file, as shown in Figure 4.

**Fig. 4.**
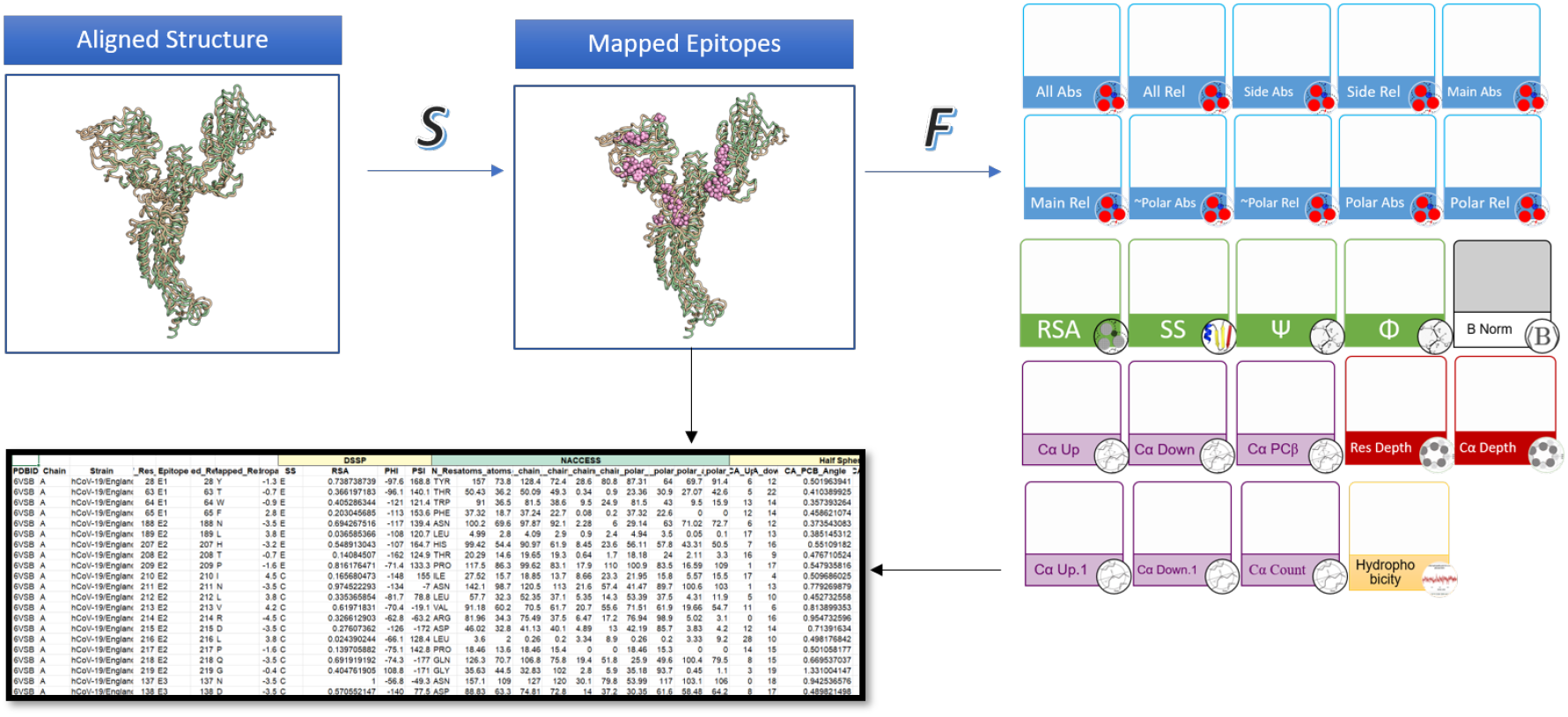
Extraction of various properties of interest from epitope residues of Spike protein. **Top left panel, ‘Aligned Structure’:** An alignment is produced in Jupytope using the StructureAlignment package of Biopython. The reference (PDB ID 7DZW, Chain A) and query (6X6P, Chain A) structures are indicated in tan and pale green ribbons, respectively. *S* is the numbered list containing epitope residue positions. **Top centre panel, ‘Mapped Epitopes’:** Mapped epitope positions are indicated as pink spheres. For clarity, only some epitopes are shown. **Right:** The angular, solvent accessible and depth related residue properties are retrieved by Jupytope as features, *F*. The properties are colored according to their source programs: NACCESS in light blue, DSSP in green, HSExposure in purple, MSMS (or residue depth) in red, the normalized B-norm (37) in grey and hydrophobicity in yellow. **Bottom left:** The mapped epitope residue positions and their corresponding structure-derived properties of interest are written to a text or csv file. One row corresponds to one epitope position. The feature properties can also be directly converted to a pandas dataframe for further work. Blue arrows represent internal stages of information flow within the program, while black arrows symbolize final output visible to the user. Up to 25 features can be extracted by the program (see Appendix for full list of features)

A total of 19,937 and 80,484 rows of raw data were extracted for the epitope residues of Influenza HA and SARS-CoV2 Spike proteins, respectively. Rows for which information was missing (for instance, when there was no corresponding query residue in the alignment for a given position) were deleted. Columns such as B-factors were also deleted when not meaningful – e.g., for spike crystal structures which were obtained from cryo-EM.

## Results and Discussion

A total of 19,922 and 54,484 cleaned rows of data were obtained for the epitope residues of Influenza HA and SARS-CoV2 Spike proteins, respectively and referred here as ‘structure-derived epitope features’. The data are are available for download from the Github at https://github.com/shamimarashid/Jupytope, and form the basis for the experiments and findings below. Although epitope prediction is not the focus of this manuscript, the extracted data are well suited as a positive dataset for training any machine or deep learning algorithm. A negative dataset may just as easily generated by supplying an appropriate list of ‘non-epitope’ residue numbers.

### Structure-derived properties of Viral Epitopes

From an initial set of 25 features (see Appendix), 16 distinct features were obtained after some overlapping, non-informative features were removed, using Pearson’s correlation analysis. Examples of removed features include internal correlations within NACCESS (absolute main or side chain solvent accessibilities and external correlations such as with DSSP solvent accessibility). The resultant epitope structural characteristics for the H1N1 and H3N2 subtypes of Influenza were compared in a distribution plot, given in Figure 5.

**Fig. 5.**
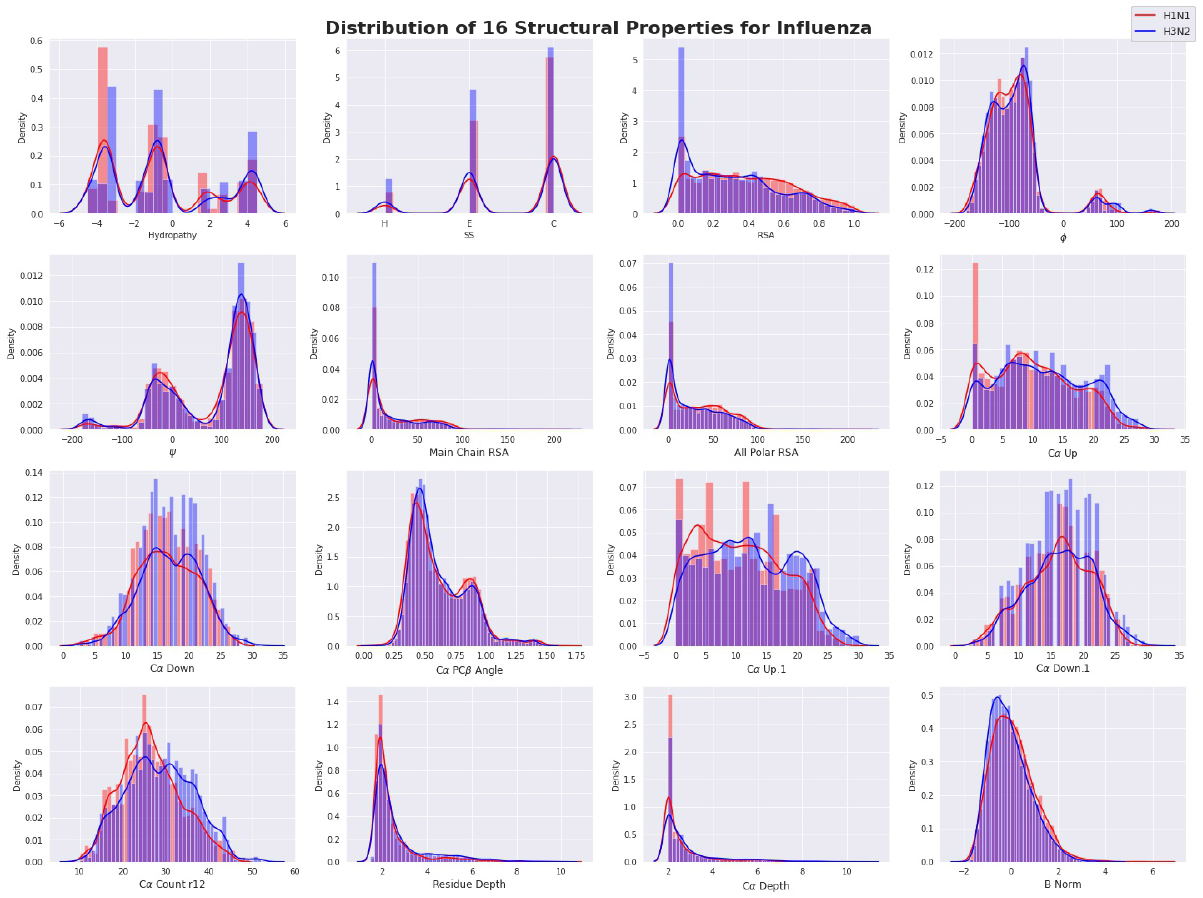
Distribution of 16 Structural Properties of Influenza from data extracted with Jupytope. H3N2 is indicated in blue, H1N1 in red and overlapping regions appear purple.

Many of the features, show a similar trendline or distribution for both H1N1 and H3N2 subtypes. Particularly, the peaks of the three secondary structure types (H = Helix, E =Sheet and C = Coil + Others) are identical for both subtypes, showing that coil structures are almost 6 times as likely to occur in the epitope residues compared to Helices. Similarly, the distribution patterns of the two subtypes are close for the relative solvent accessibility (RSA) and *ϕ*, *Ψ* dihedral angles with only a small difference occurring near the peaks. While visually these features do not seem informative in distinguishing the subtypes, they are shown to contribute to the XGBoost model performance and gains (described below).

However, a stronger difference seems to be observed for the features involving C*α* atoms CA Up and CA Down (corresponding to HSE*α*) as well as CA Up.1 and CA Down.1 (corresponding to HSE*β*). The most interesting difference seems to be for CA Count r12, which is the no of surrounding C*α* atoms within a 12*Å* sphere of the epitope residue. The visible difference in distribution peak is minor for B_Norm, so it is not expected to be that informative.

To quantitatively rank the feature importance and to critically examine their role in viral subtype or clade detection, a simple XGBoost classifier is modeled on the Spike and HA datasets.

### Application of structure-derived epitope features for Clade and Subtype Detection – Demonstration with XGBoost

To assess if the extracted epitope data are associated with virus groups (i.e. clade or subtype), the features are compared to a random labelled dataset which serves as a baseline. It was hypothesized that if the epitope properties carried antigenic profiles specific to virus groups, the classification model would show an improvement against a random baseline. Otherwise, if the structure-derived epitope features simply encoded generic residue properties, the model’s performance would be comparable for both true and random labels.

The alphabetical clade or subtypes (i.e. groups) were first converted to numeric class labels as follows: Subtype = {H1N1*→*0, H3N2→1} and Clade = {G→0, L→1, O→2, S→3, V→4} for Influenza and SARS-CoV2, respectively. Clade G includes all of the sub-clades (such as GH, GRY, GK, etc.) The random labels were assigned by generating random integers as ‘psuedo’ numerical labels.

The distribution of epitope residues with the actual (or original) labels) and randomly assigned labels for each virus group are shown in Figures 6a and 6b below. Clearly, the random labels follow an equal distribution while the actual labels are distributed unevenly between virus groups.

**Fig. 6.**
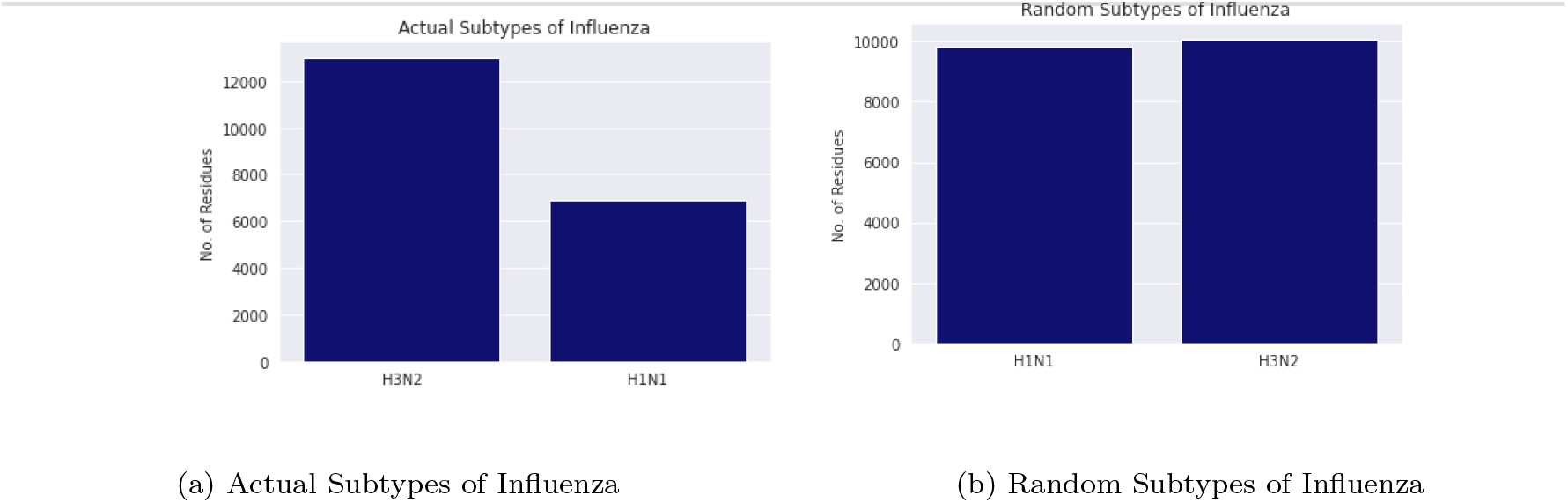
Distribution of Influenza Labels by Subtype

An XGBoost model (xgboost version 1.0.2 of scipy library, version 1.5.4) was trained on the Influenza and SARS-CoV2 datasets for both actual and random virus labels. The data was split with 75% used as training and 25% as test samples. The extracted structure-derived epitope properties were the model’s features.

All parameters were kept as default and no attempts were made to search for better hyperparameters, except the random state which was set to 1. This allows reproducibility of results and facilitates comparison across different virus types. The findings also establish the minimum bar of performance for associations between epitope properties and virus groups present in the data. Such structure-derived epitope-group associations cannot be attributed to fluctuations in randomized model states or one-off stellar outcomes from user-defined optimization of model parameters with respect to the dataset.

Since clade ‘O’ is default for any strain that does not belong in {G, L, S, V}, it may contain strains that are not specific to any particular lineages or lack antigenic profile signals that are typical of any defined SARS-CoV2 clade. Hence data rows labelled as group ‘O’ were removed and and the XGBoost results were also computed for a 4-clade classification problem. The findings from using different numbers of features (23, 15 and 10), for both 4 and 5-Clade classification problems, are presented in Figure 8.

**Fig. 7.**
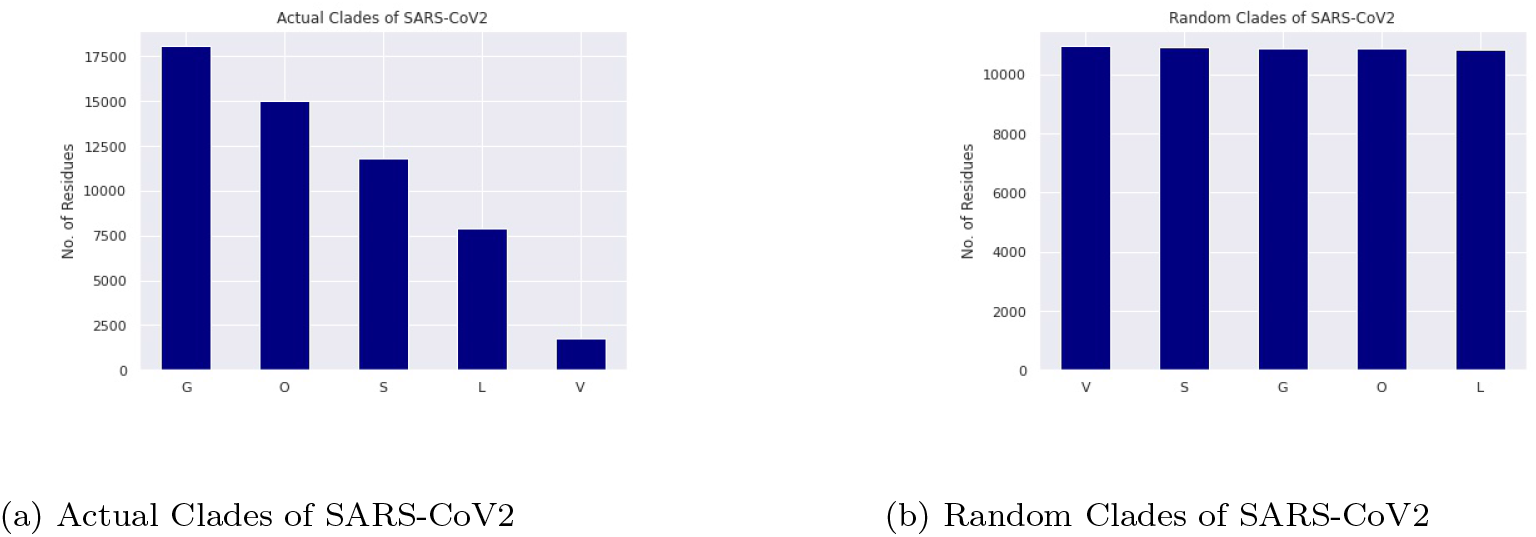
Distribution of SARS-CoV2 Labels by Subtype

**Fig. 8.**
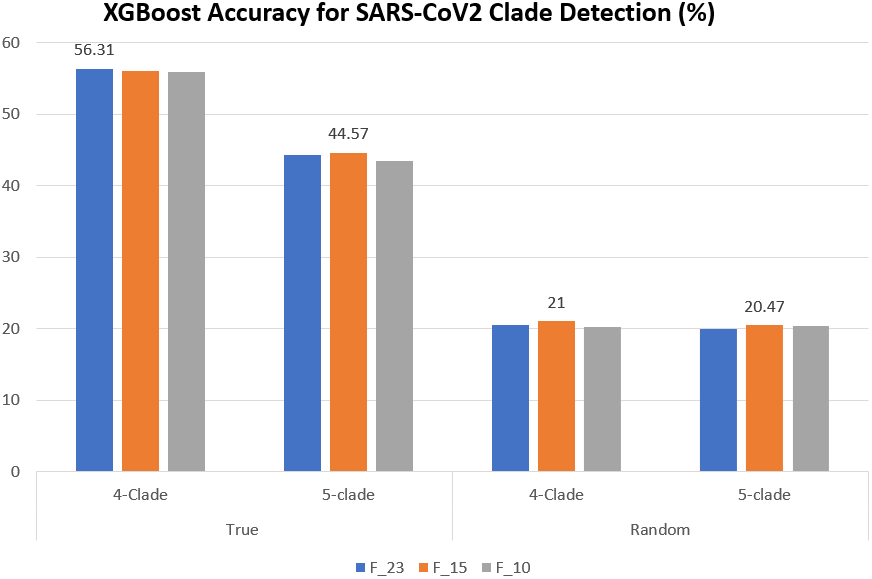
Accuracy of XGBoost Model, SARS-CoV2. Bars are grouped according to the classification type. 4-clade classification predicts the following major clades: G, L, S and V while 5-clade includes clade group ‘O’ in the prediction. True and Random represent the type of labels presented to the model. Colours indicate the number of features used - 23(blue), 15(orange) or 10(grey). 23_F includes all the features except CA_CB angle and B_Norm (see Appendix). The highest accuracy in each type of prediction is indicated above the respective bar.

For the 5-Clade classification, the highest accuracy obtained was 44.6% using 23 features for actual clades as labels, compared to 20.5% using random labels. The accuracy of random labels is as expected from pure chance (maximum entropy), equivalent to a fifth of all outcomes. Even though the figure of 44.6% shows much room for improvement, the features contribute twice the information compared to chance alone.

The best accuracy for SARS-CoV2 clade detection was 56.3% for true clades, obtained using 23 features for the 4-Clade classification. We also tested the 15 features in Figure 5 for Influenza (excluding B Norm), but the accuracy was slightly lower at 56.1%.

For SARS-CoV2, regardless of the number of clades (4 or 5) or the number of features used, the accuracy for random labels remains close to 20%. The strong improvement of true labels compared to random labels clearly indicates the usefulness of these features for clade detection and indicates the presence of an antigenic profile that is associated to clades. For Influenza, the findings are even stronger with the XGBoost model obtaining an accuracy of 93.2% which is a large improvement of 43.4% over the random clade labels (shown in Figure 9). The strong improvement of true labels compared to random labels, and the excellent accuracy for Influenza, indicates the usefulness of these features for clade or subtype associations.

**Fig. 9.**
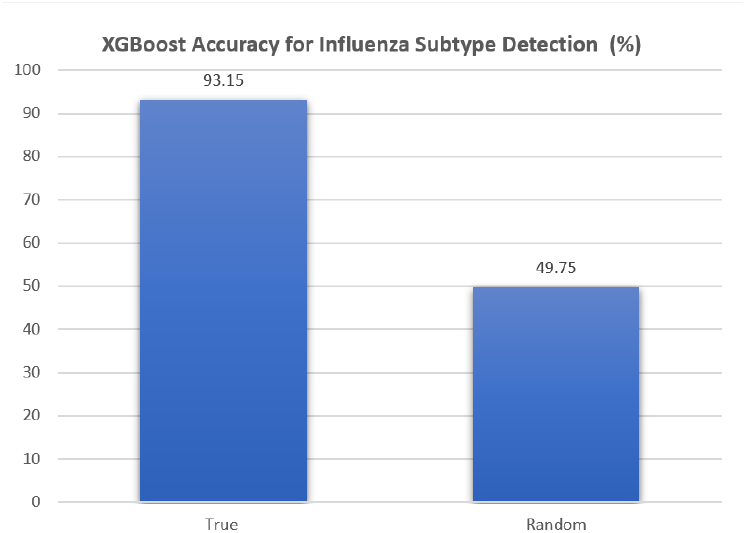
Accuracy of XGBoost model, Influenza. The subtypes considered are H1N1 and H3N2. True and Random represent the type of labels presented to the model.

To verify the reliability of the XGBoost results, 100 runs for each experiment type were conducted and reported in Table 2. The randomized state was allowed to vary with each model instantiation, and a fresh train to test split (at 75% and 25% respectively) was taken at the beginning of each run. Table 2 indicates that results for true labels are very similar to those in Figures 9 and 8 with average accuracies differing by less than 1%. Standard deviations (s.d.) are given in brackets in Table2. These are very small (<0.001) and indicate that the findings are stable and repeatable across the model’s trials.

**Table 2.** Average Best Accuracy of 100 runs

| Virus | True % (s.d.) | Random % (s.d.) |
| --- | --- | --- |
| Influenza | 92.57 (0.004) | 50.14 (0.006) |
| SARS-CoV2 | 56.35 (0.004) | 24.91 (0.003) |

The feature gains in XGBoost describe features that contribute to the model’s internal abstractions of what features are contributive or associative to the five clades or the subtypes H1N1 or H3N2. Such features serve as proxies for the association of structure descriptors with clades or subtypes. The highest feature gains contribute the most to model performance, and are of interest. Figure 10 indicates the top gains for SARS-CoV2 and Influenza datasets, respectively. The gains for true labels (clade or subtype, respectively) are compared with those of random labels. Figures 10a and 10b suggest Hydropathy and SS for both virus groups, with Influenza showing a substantial gain over the random labels.

**Fig. 10.**
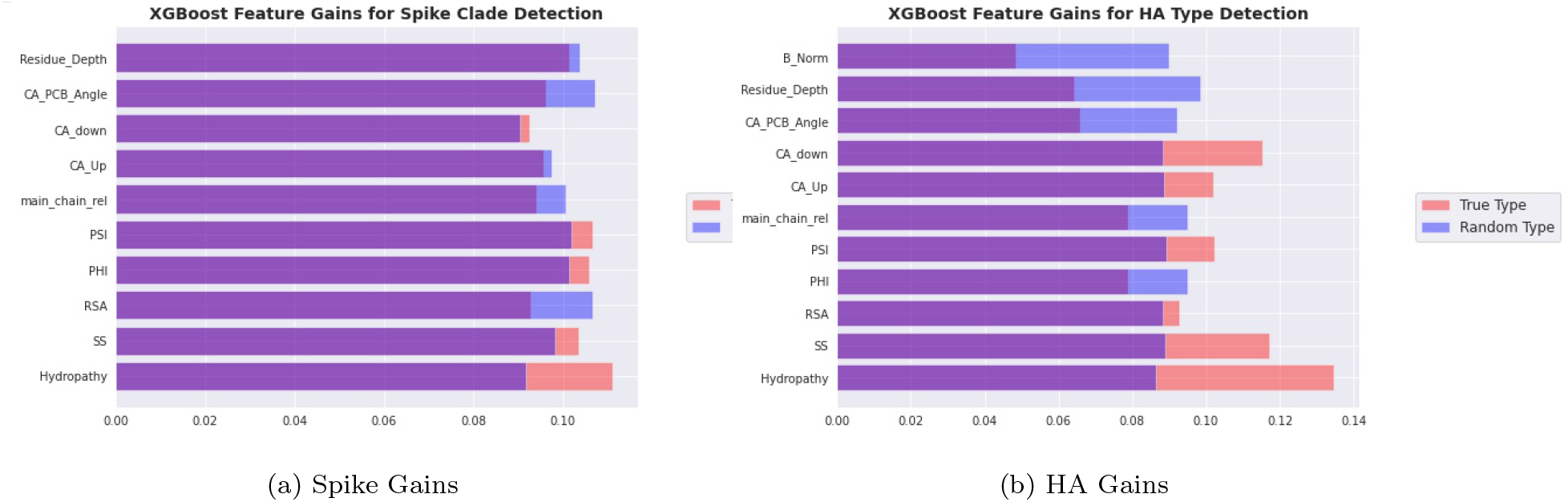
Top Feature gains identified by XGBoost. True clade and true type refer to the actual clade and subtype assignments, while ‘Random’ indicates random labels. Feature B_Norm is excluded for Spike clade detection since it contains only cryo-electronmicrography (cryo-EM) crystals, where B-factors may not be applicable.

The CA down, which is a HSE measure for the number of C*α* atoms in the lower sphere of the plane, is also a significant contributor to the high accuracy achieved in Influenza subtype classification.

Surprisingly, while the feature distribution in Figure 5 had suggested that atom environment descriptors like CA_up or C_down showed more distinct patterns between H1N1 and H3N2, the biggest model gains were in fact from ‘simpler’ descriptors such as secondary structure (SS). Conversely, residue depth and B-factors appeared to be ‘generic’ protein features, with equal or greater associations with random labels.

The resolution of crystal structures could have been a factor in the lower accuracy for SARS-CoV2 that affected the model’s performance. It suggests the need of algorithms capable of handling low resolution PDB structures for prediction, to maximize the available data for training.

## Conclusion

This work presents a tool to quickly extract structure-derived properties from viral epitopes of interest, that can be conveniently saved in text or csv formats. Alignment to a user-defined reference structure ensures correct mapping and preservation of spatial characteristics of epitopes.

Importantly, the structure-derived epitope features were shown to recover clade or subtype-specific associations in SARS-CoV2 Spike and Influenza HA proteins, respectively. Performance of a simple XGBoost model in classifying SARS-CoV2 clades or Influenza subtypes showed much better performance compared to a random baseline, especially for Influenza. The findings indicate that the extracted properties are not generic solvation or structural descriptors of an epitope residue’s 3-dimensional environment, but that they can capture chemical and atom environment profiles related to antigenic characteristics. The work also provides a dataset of SARS-CoV2 PDB structures annotated with Clade and Pango lineages. Overall, the tool and annotated datasets are expected to aid structural studies or predictive work on viral epitopes. The extracted epitope data can be used in conjunction with sequence alignment models to construct antigenic profiles in future works.

## List of Extracted Structural Properties

1. Hydropathy

### DSSP

2. SS - Secondary Structure, DSSP
3. RSA – Relative Solvent Accessibility
4. PHI – Dihedral angle
5. PSI – Dihedral Angle

### NACCESS Solvent Accessibilities

6. all_atoms_ abs – All Atom Absolute
7. all_atoms_ rel – All Atom Relative
8. side_chain_abs – Side Chain Absolute
9. side_chain_rel – Side Chain Relative
10. main_chain_abs – Main Chain Absolute
11. main_chain_rel – Main Chain Relative
12. non_polar_abs – Non Polar Absolute
13. non_polar_rel – Non Polar Relative
14. all_polar_abs – All Polar Absolute
15. all_polar_rel – All Polar Relative

### Half Sphere Solvent Exposure (HSE)

16. CA_Up - C*α* Up, HSE*α*
17. CA_down - C*α* Up, HSE*α*
18. CA_PCB_Angle - C*α* -PC*β* Angle, HSE*α*
19. CA_Up.1 - C*α* Up, HSE*β*
20. CA_down.1 - C*α* Down, HSE*β*
21. CA_CB_Angle - C*α* -C*β* Angle, HSE*β*
22. CA_Count_r12 -C*α*, within 12*Å* radius

### MSMS

23. Residue_Depth – Residue Depth
24. CA_Depth – C*α* Depth
25. B_Norm – Normalized B Factor

## Acknowledgements

This work was funded by Ministry of Education (MOE) grants MOE2019-T2-2-175 and MOE2020-T1-001-130, Singapore. The authors thank Jason Brownlee of machinelearningmastery.com for concise code of an XGBoost model that was used here with modifications

**Shamima Rashid** holds a PhD, and is a Research Fellow in the Biomedical Informatics Lab in the School of Computer Science and Engineering, Nanyang Technological University. Her research interests include protein structural bioinformatics and machine learning.

**Ng Teng Ann** holds a B.Eng. in Computer Science and is currently working as a Project Officer in the research scheme at the School of Computer Science and Engineering, Nanyang Technological University

**Chee Keong Kwoh** holds a PhD Degree from the Imperial College of Science, Technology and Medicine, University of London. He is currently an Associate Professor of the School of Computer Science and Engineering, Nanyang Technological University (NTU). His research interests include data mining, soft computing and graph-based inference, and applications areas include bioinformatics and biomedical engineering.

